# A Low-Cost, High-Throughput Design-Build-Test Pipeline for Engineering Genetic Systems: Stress Testing with Complex Structural Proteins

**DOI:** 10.64898/2026.06.08.729977

**Authors:** Harry E. Adamson, James R. McLellan, Khushank Singhal, Melik C. Demirel, Howard M. Salis

## Abstract

Genetic systems engineering is constrained by high DNA synthesis costs, assembly inefficiencies, and challenges in expressing complex proteins. To address these limitations, we developed a highly parallel, low-cost pipeline for the design, assembly, and functional screening of genetic systems, which we stress-tested on highly repetitive structural proteins, including spider silk, biocements, reflectins, and talins. The integrated pipeline combines computational genetic systems design, low-cost many-plasmid DNA assembly from oligopools, automated many-to-many mapping using nanopore sequencing data, and a label-free biosensor to measure single-cell protein expression levels. We applied this pipeline to build 240 plasmids, achieving an 88% success rate (up to 2000 bp) using standard clonal isolation and 58% assembly efficiency (up to 5600 bp) without selective DNA purification, while lowering material costs by up to 24-fold. We applied the biosensor to identify genetic factors that create distinct cellular subpopulations with varying protein expression levels. Overall, the integrated pipeline will dramatically lower the cost of high-throughput synthetic biology, while demonstrating how designing genetic systems to improve build efficiency (“design for build”) and directly incorporating biosensors into genetic systems (“design for test”) will greatly accelerate design-build-test workflows.

## INTRODUCTION

The Design-Build-Test (DBT) cycle drives synthetic biology applications, but scaling genetic engineering faces critical bottlenecks in sequence design, DNA synthesis & assembly, and functional screening^1^. Advances in generative AI can design large sequence libraries, yet experimentally validating these constructs remains challenging, especially for repetitive or GC-rich structural proteins prone to synthesis, cloning, and expression errors^2-5^. Overcoming these limitations demands an integrated DBT approach that streamlines genetic design, improves cloning efficiency, and enhances protein expression screening, while lowering costs.

A key challenge in genetic design is that genetic parts cannot be combined in a predictable manner without taking into account the entire genetic system sequence. When inserted into a new genetic system, even well-characterized genetic parts can introduce undesired interactions that take place between and within genetic parts^6^. For example, combining genetic parts can create repetitive DNA sequences that trigger homologous recombination, leading to DNA mutation and deletion^7^. The presence of regulatory sequence motifs (e.g. promoters, ribosome binding sites, terminators) inside other genetic parts (e.g. coding sequences) can lead to aberrant expression of undesired RNAs and proteins as well as confounding interactions that can interfere with the desired functions^8, 9^. Model-based design algorithms can calculate how genetic part sequences control transcription, translation, and mRNA decay rates^10-14^, which is essential to achieving desired expression levels without laborious screening. However, multi-objective design algorithms are needed to achieve desired expression levels, while eliminating undesired sequence motifs that can break the DBT cycle. Multi-objective design algorithms can remove these failure modes by designing genetic part sequences and many-part system sequences to simultaneously satisfy several design criteria. Here, we combine the Promoter Calculator, RBS Calculator, and CDS Calculator to carry out multi-objective design of genetic systems to control expression levels, while eliminating undesired sequence motifs across the entire genetic system (**Figure 1A**).

**Figure. 1.**
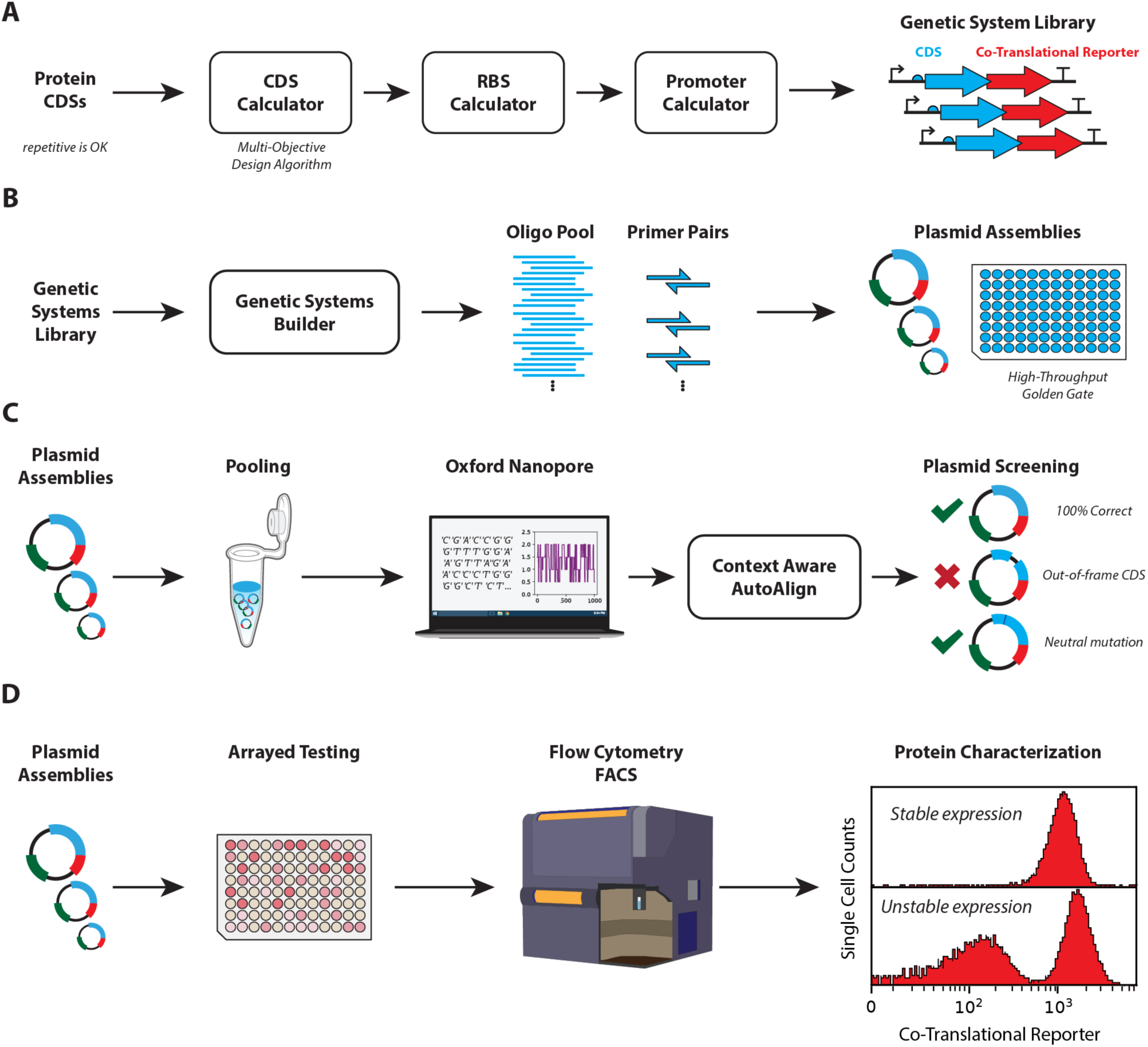
High-throughput, parallelized design-build-test pipeline for engineering genetic systems. **A** An automated process combines multiple models (Promoter Calculator, RBS Calculator, CDS Calculator) to design promoters, ribosome binding sites, and protein coding sequences. Several design rules are applied to remove undesirable sequences, including repetitive nucleotide sequences, internal promoters, internal translated open reading frames, and pause motifs, while achieving targeted transcription and translation rates. Protein coding sequences are designed to be co-expressed with a co-translational reporter protein that produces two separate proteins. **B** The Genetic Systems Builder designs the oligonucleotides and PCR primers to build many plasmids from a single oligopool. The algorithm uses dynamic programming to identify optimal overhangs within each genetic system sequence for optimal Golden Gate assemblies, followed by the design of high-specificity PCR primers. **C** Assembled plasmids are pooled and sequenced together using long-read nanopore sequencing, followed by mapping to all reference sequences using the Auto Align algorithm. **D** The distribution of single-cell protein expression levels is measured using flow cytometry, leveraging a dye-free, label-free co-translational reporter that measures the expression of the protein of interest. Measuring single-cell expression enables the identification of heterogeneity in a cell population, due to genetic instabilities or other environmental factors.

Building many genetic systems remains a slow, costly process that often fails for “complex” sequences^3, 15^. Constructing even a modest library of genetic systems at current commercial rates becomes prohibitively expensive. For example, at the current commercial prices for non-clonal DNA fragments ($0.07 to $0.09 / bp) or clonal, sequence-verified DNA fragments ($0.12 to $0.25 / bp)^16^, building 100 different genetic systems (3000 bp inserts) costs anywhere from $20000 to $75000, depending on the assembly efficiency and the number of sequenced clones. However, beyond cost, many genetic system sequences are too “complex” to be constructed correctly. Whenever a genetic system sequence contains >75% GC regions or long repetitive sequences, there is a higher chance of mis-annealing or mis-priming during DNA assembly, leading to large DNA deletions^17^. In particular, genetic systems that express structural proteins (e.g. spider silks and biocements) are notorious for having such “extremely complex” sequences (e.g. many tandem repeat sequences) that commercial DNA synthesis providers will outright refuse to build them.

The recent development of precision oligopool synthesis has provided a route to overcoming these build challenges. Oligopools are complex mixtures of DNA oligonucleotides with defined sequences (150 to 350 nt, or longer) that are chemically synthesized on nanofluidic platforms with extremely high economy-of-scale (e.g. one million oligonucleotides per chip), leading to ultralow per-unit costs (e.g. $0.0004 / nt). New DNA assembly techniques are being developed to harness oligopools as low-cost DNA substrates^18-21^, while leveraging recent developments enabling many-fragment Golden Gate assemblies. For example, combining the characterization of Golden Gate overhang ligation efficiencies with algorithmic selection of optimal overhangs^22^, it is now possible to carry out one-pot Golden Gate assemblies with at least up to 52 DNA fragments^23, 24^, though the assembly efficiency highly depends on the number of DNA fragments. We and others have developed oligopool design algorithms that introduce designed PCR primer binding sites to enable multiplexed PCRs^25-28^. For example, the Oligopool Calculator rapidly designs highly selective primer binding sites for multiplexed PCRs on oligopools with up to 200 million nt of template DNA^25^. By combining primer binding site design with optimal overhang selection, it is possible to create novel DNA assembly procedures that combine PCR and many-fragment Golden Gate assembly to build many plasmids from a single low-cost oligopool.

However, maximizing the efficiency of DNA assembly depends on solving a combinatorial optimization problem subject to several design constraints. Prior efforts have utilized stochastic optimization algorithms (e.g. simulated annealing or a genetic algorithm) to propose good-enough solutions; however, these approaches can fail on complex sequences^22^. Here, we introduce the Genetic Systems Builder algorithm, which uses dynamic programming with beam search to rapidly and deterministically determine the optimal overhangs that maximize the overall efficiency of each many-fragment Golden Gate assembly, followed by designing the oligopool and primer sequences to carry out PCRs and assemblies (**Figure 1B**). We introduce a novel ligation fidelity formula that takes into account the probability that Type IIS enzymes will shift their cleavage site, producing an unexpected overhang sequence and undesired ligation reaction. The Genetic Systems Builder has several additional capabilities, including: (i) the design of circular assemblies where the many-fragment assembly includes the vector portion of the plasmid; (ii) the design of multi-level DNA assemblies, utilizing multiple Type IIS enzymes and assembly junctions, to build even longer constructs; and (iii) the design of genetic system libraries where many module variants can be combined together using combinatorial assembly.

Long-read nanopore sequencing is often the preferred approach for confirming DNA assembly correctness. However, analyzing long read data across hundreds of assembled plasmids remains laborious. To accelerate this step, we introduce Auto Align, a long-read mapper that automatically identifies correct plasmids from a long list of expected sequences, using either mixed or consensus long reads (**Figure 1C**). Whenever mutations are identified, the algorithm predicts changes in bacterial expression levels (a change in transcription or translation rate) as well as any change in the protein’s amino acid sequence, enabling one to determine the significance of the mutations.

Finally, measuring protein expression levels remains a major bottleneck in the DBT workflow. While bulk or lytic techniques can be scaled-up in microtiter plates, they obscure non-clonal, heterogenous phenotypes by only measuring population-averaged protein levels^29-31^. Modifying proteins with fluorescent or luminescent tags can also disrupt protein folding and function^32^. Here, we demonstrate a non-tagged, label-free, single-cell fluorescent biosensor that harnesses translational coupling to express a fluorescent protein in direct proportion to the protein of interest, leveraging our Translational Coupling Calculator^13^ to ensure correct design of the ribosome reinitiation site (**Figure 1D**). We then carry out flow cytometry on non-lysed cells to quantify protein expression levels with single-cell resolution. The resulting fluorescence distribution reveals any hidden population heterogeneity, providing the critical information needed to measure the functional differences across genetic system designs.

We stress-tested this integrated DBT pipeline on a large set of heterologous structural proteins that are particularly challenging to express in *E. coli* cells. We first selected 16 structural proteins (**Table 1**), including spider silks, biocements, and reflectins, and built designed RBS libraries to optimize their protein expression levels. We then scaled-up the pipeline to build 240 plasmids, including circular assembly of vectors. Overall, the integrated DBT pipeline achieved a 78% assembly efficiency with an over 10-fold lower material cost per plasmid, while identifying engineered strains that achieved both over-expression and genetic stability. With these concrete examples, we show how essential engineering principles – “design for build” and “design for test” – will accelerate the design, assembly, and screening of complex genetic systems.

**Table 1.**
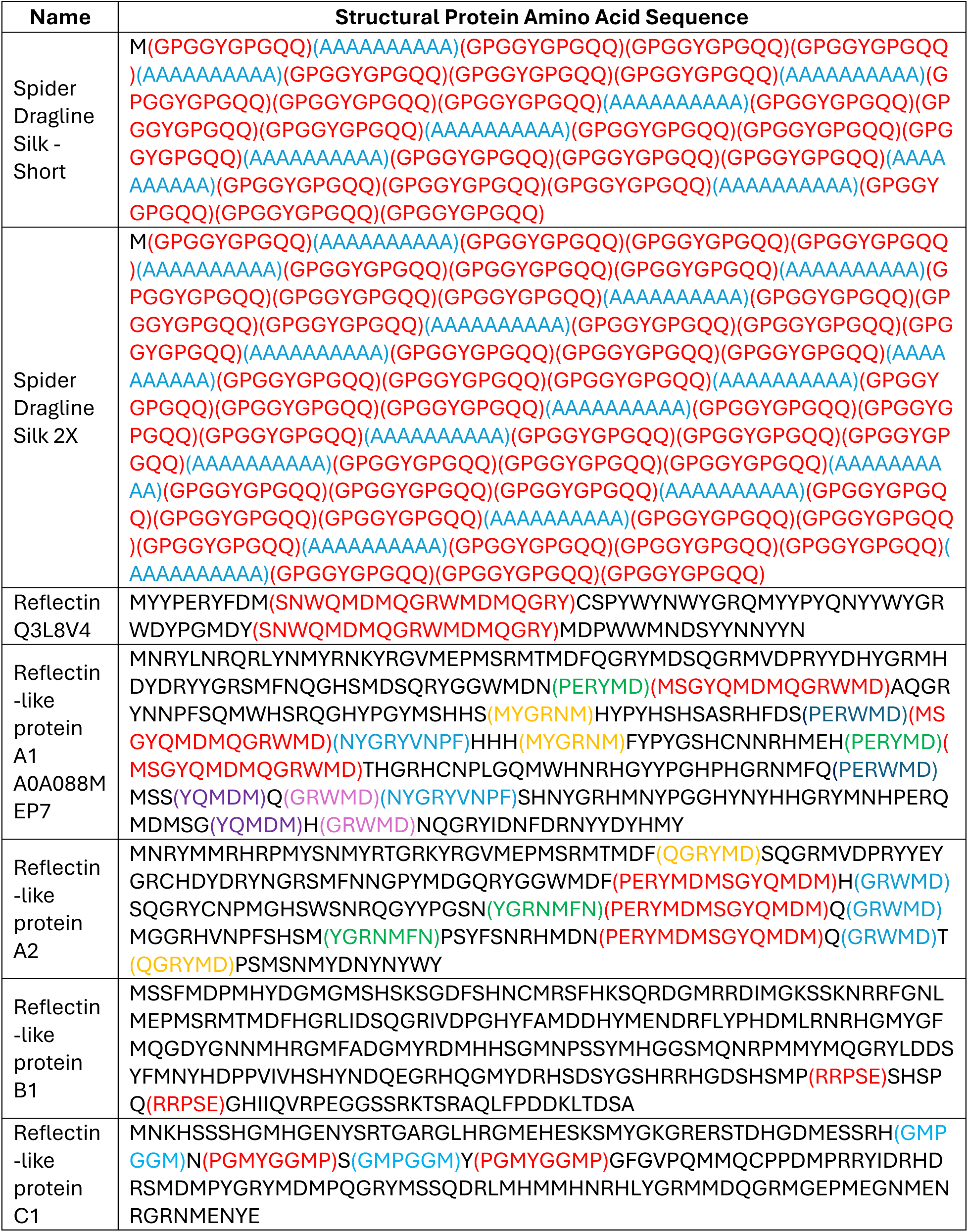
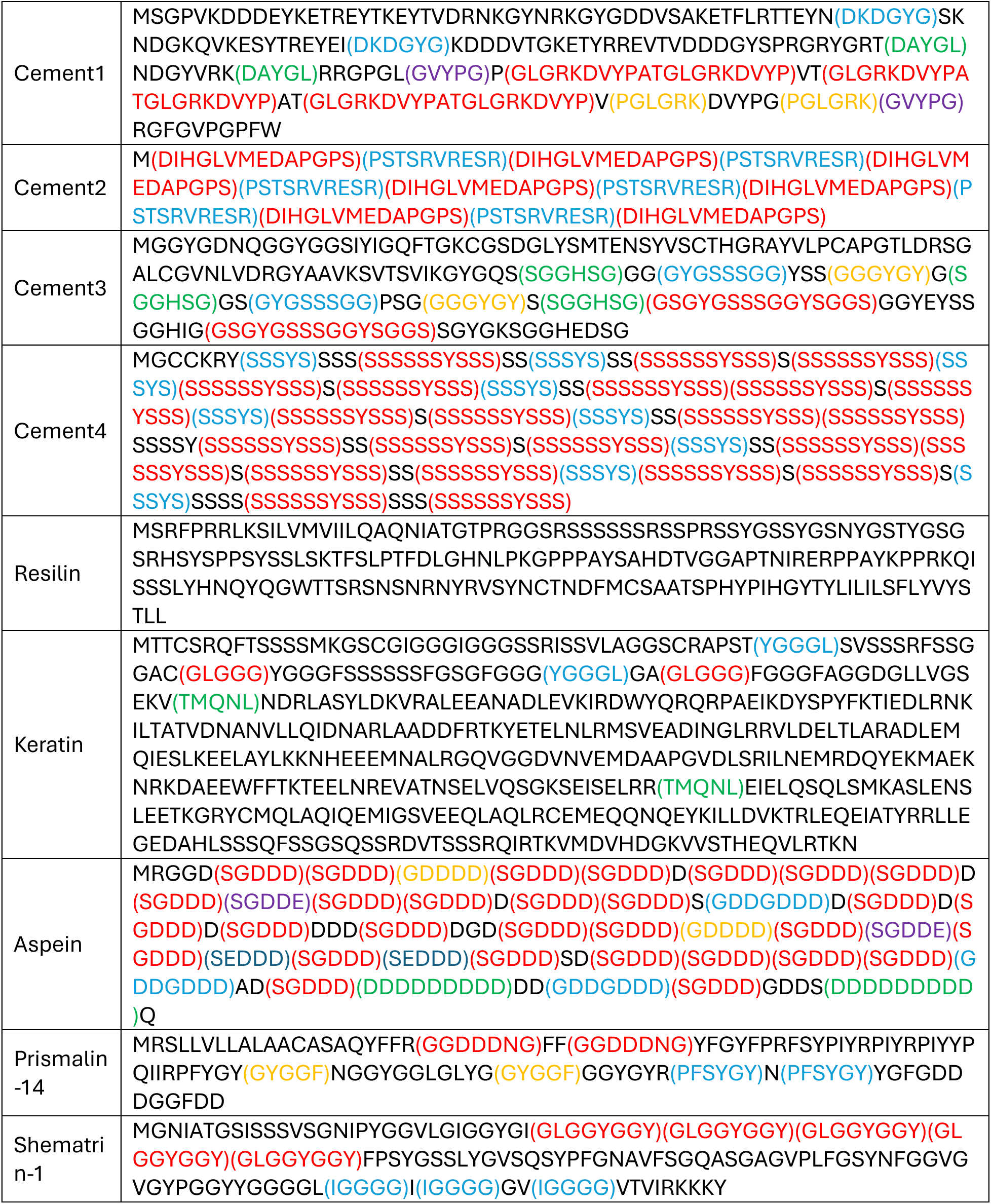
Structural protein sequences. Protein name and amino acid sequence are shown for each structural protein used to test the DBT pipeline.

## RESULTS

### A Design Pipeline for Protein Coding Sequence Optimization

The pipeline begins by redesigning protein coding sequences using the CDS Calculator optimization algorithm. The CDS Calculator carries out multi-objective synonymous codon optimization with the overall goal of maximizing predictable control over the protein’s expression level, including reducing the production of undesired mRNA or protein isoforms, removing repetitive DNA sequences to improve genetic stability, removing internal transcriptional terminators and ribosomal pause sites, and modifying codon usage to increase translation elongation rates (**Figure 2A**). The algorithm uses adaptive path generation to step through the synonymous codon choice decision tree, using our list of model predictors and adaptive design rules to determine sequence correctness. Each step through the decision tree is a random choice of synonymous codon, weighted by the genome-specific synonymous codon frequency table. After each step, the algorithm sequentially evaluates the correctness of the partially generated CDS sequence according to an extensible list of design rules. Each design rule returns a binary decision (success or failure) along with the beginning position of the sequence that caused the design rule failure. If all selected rules succeed, the path generation continues. If any selected rule fails, the algorithm rewinds the state of the decision tree to the sequence position where the rule failure was introduced. The path generation process is stochastic, enabling exploration of synonymous codon choices as paths are generated. If path generation succeeds with the final choice, the fully generated CDS sequence is returned. This approach is similar in concept to our previous Non-Repetitive Parts Calculator algorithm^7^; here, we expand the concept of model-based constraints to include any type of design rule.

**Figure 2.**
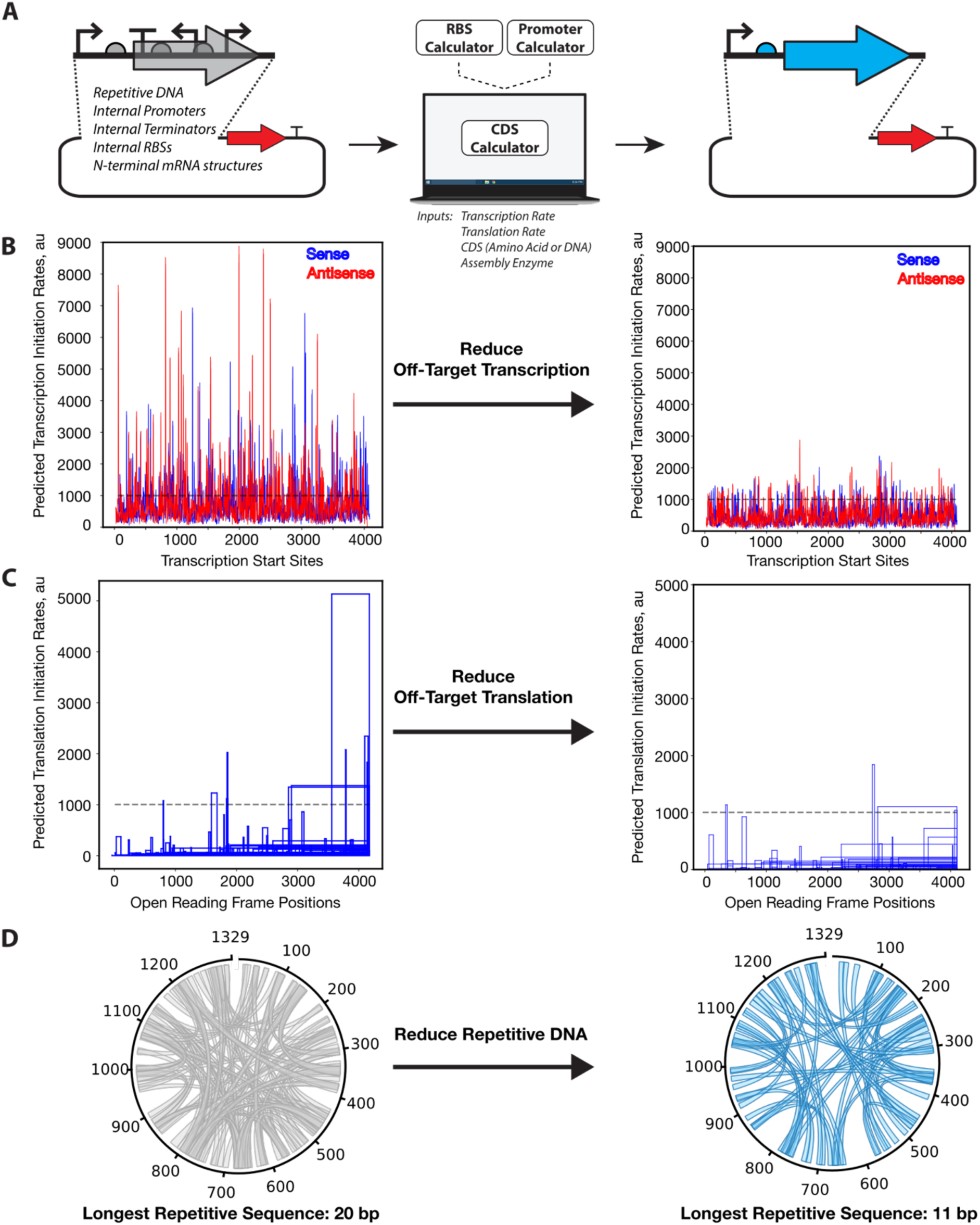
CDS optimization examples. **A** Schematic showing how the CDS Calculator redesigns protein coding sequences to control expression and remove undesired sequences. **B** Predicted transcription initiation rates across the (left) natural Cas9_SP_ CDS sequence and the (right) Cas9_SP_ CDS sequence designed by the CDS Calculator. The adaptive design rule for internal transcription uses an initial threshold of 1000 au (black dotted line). **C** Predicted translation initiation rates of the internal open reading frames of the (left) natural Cas9_SP_ CDS sequence and the (right) Cas9_SP_ CDS designed by the CDS Calculator. **D** Repeat chord diagram showing the location of repetitive DNA motifs inside the (left) natural squid ring teeth protein sequence and the (right) squid ring teeth protein sequence designed by the CDS Calculator. The longest repetitive DNA sequences are shown.

Whenever many design rules are applied simultaneously using constant criteria, there is a frequent possibility that overlapping sequence-function constraints eliminate all possible sequence solutions. To overcome this challenge, we introduce adaptive design rules that enable the CDS Calculator to return a nearly optimal CDS sequence in a finite number of iterations. To do this, the algorithm tracks the number of rule failures at each nucleotide position in the CDS sequence, which we call “design complexity”. Every design rule takes into account the position-specific design complexity when determining rule success or failure. For example, to eliminate internal promoters inside the protein CDS, we first apply the Promoter Calculator model to predict transcription initiation rates at start sites within the partially generated CDS sequence. We define a design rule that returns a failure whenever the predicted transcription initiation rate exceeds a threshold. Without any design complexity, the threshold ***Y*** begins at 1000 au on the Promoter Calculator scale. Whenever the design complexity ***X*** exceeds a minimum failure threshold ***F***, we then increase the threshold by an amount ***dY*** = ***M*** * (***X*** – ***F***), where ***M*** is a defined rate of rule relaxation. Repeated failure of the “remove internal promoters” design rule at the same nucleotide position will incrementally increase the rule’s threshold, enabling the path generation algorithm to efficiently search for CDS sequences until it finds one that has a nearly minimal amount of internal transcription rate. Overall, we define 15 adaptive design rules to carry out CDS sequence design.

We first demonstrate the CDS Calculator on two example CDSs expressing Cas9 from *S. pyogenes* and squid ring teeth (SRT) protein^33^. The natural CDS sequence for Cas9 contains 224 transcriptional start sites predicted to have moderate to high transcription initiation rates (3000 to 8900 au in *E. coli*) that can produce inhibitory antisense RNA as well as truncated mRNA transcripts (**Figure 2B**). Similarly, the natural Cas9 CDS sequence is predicted to have 4 open reading frames with moderately high translation initiation rates (2000 to 5100 au in *E. coli*) that can produce in-frame, truncated proteins and out-of-frame, distinct proteins (**Figure 2C**). After redesigning it with the CDS Calculator, the Cas9 CDS sequence did not have any predicted transcriptional start sites with transcription initiation rates over 3000 au, while it had no internal open reading frames with translation initiation rates over 2000 au (**Figure 2BC**). To carry out the redesign process, the CDS Calculator modified 817 synonymous codons out of Cas9’s total of 1369 codons. As a structural protein, the natural ring teeth protein has a highly repetitive amino acid and nucleotide sequence with a maximum repeat length of 20 bp (**Figure 2D**), which is sufficiently long to trigger homologous recombination and genetic deletions. After redesigning it, the CDS Calculator identified synonymous codons that greatly reduced the repetitiveness of the nucleotide sequence, while maintaining the amino acid sequence, with an overall maximum repeat length of only 11 bp (**Figure 2D**). For comparison, the common J23100 promoter has a predicted transcriptional initiation rate of about 10000 au on the Promoter Calculator’s scale (depending on sequences upstream and downstream of the core promoter).

After the CDS sequence is designed, we use the RBS Calculator to design a synthetic ribosome binding site sequence with a target translation initiation rate. In some scenarios, we then use the Promoter Calculator to design a synthetic constitutive promoter sequence with a target transcription initiation rate. Alternatively, other types of promoters may be selected, for example, engineered T7 promoters that utilize lacO sites to enable IPTG induction. To date, over 590 researchers have applied the CDS Calculator to design over 7200 coding sequences that express a wide variety of proteins, utilizing a web-based interface at https://salislab.net/software. Since 2021, over 3500 researchers have utilized the Promoter Calculator model on over 110000 DNA sequences. Since 2019, over 10000 researchers have utilized the RBS Calculator model on over 632000 mRNA sequences.

### Design of Complex Structural Protein Coding Sequences

We applied the CDS Calculator to design the protein coding sequences of 16 structural proteins that have complex sequences, including highly repetitive amino acid motifs and highly biased amino acid usage. For example, spider dragline silk 2X contains 49 repeats of a GPGGYGPGQQ elastomer spring-like motif and 16 repeats of a AAAAAAAAAA rigidity motif, the shell matrix protein (aspein) from pearl oysters contains 28 repeats of a mineral-binding SGDDD motif, and an aquatic calcium-binding biocement protein has 279 serine residues (over 87% of its length).

Overall, the CDS Calculator designed protein coding sequences with minimally low rates of internal transcription and minimally low translation rates of internal ORFs, while minimizing the repetitiveness of the DNA sequences (**Figure 3**). Across all the 16 structural protein CDSs, there were zero transcriptional start sites with even moderate levels of predicted transcriptional activity (3000 au or higher) and most had maximum predicted transcription initiation rates below 2000 au (**Figure 3A**). Almost all of the structural protein CDSs had low internal translation rates (around 1000 au or less), except for the aspein protein CDS, which contains numerous AGG and ATG-rich motifs that cause high internal translation rates (**Figure 3B**). All of the structural protein CDSs had minimally repetitive DNA sequences (the maximum length of repeats was shorter than 20 bp), except for the 2X version of spider dragline silk protein and the aspein protein (**Figure 3C**).

**Figure 3.**
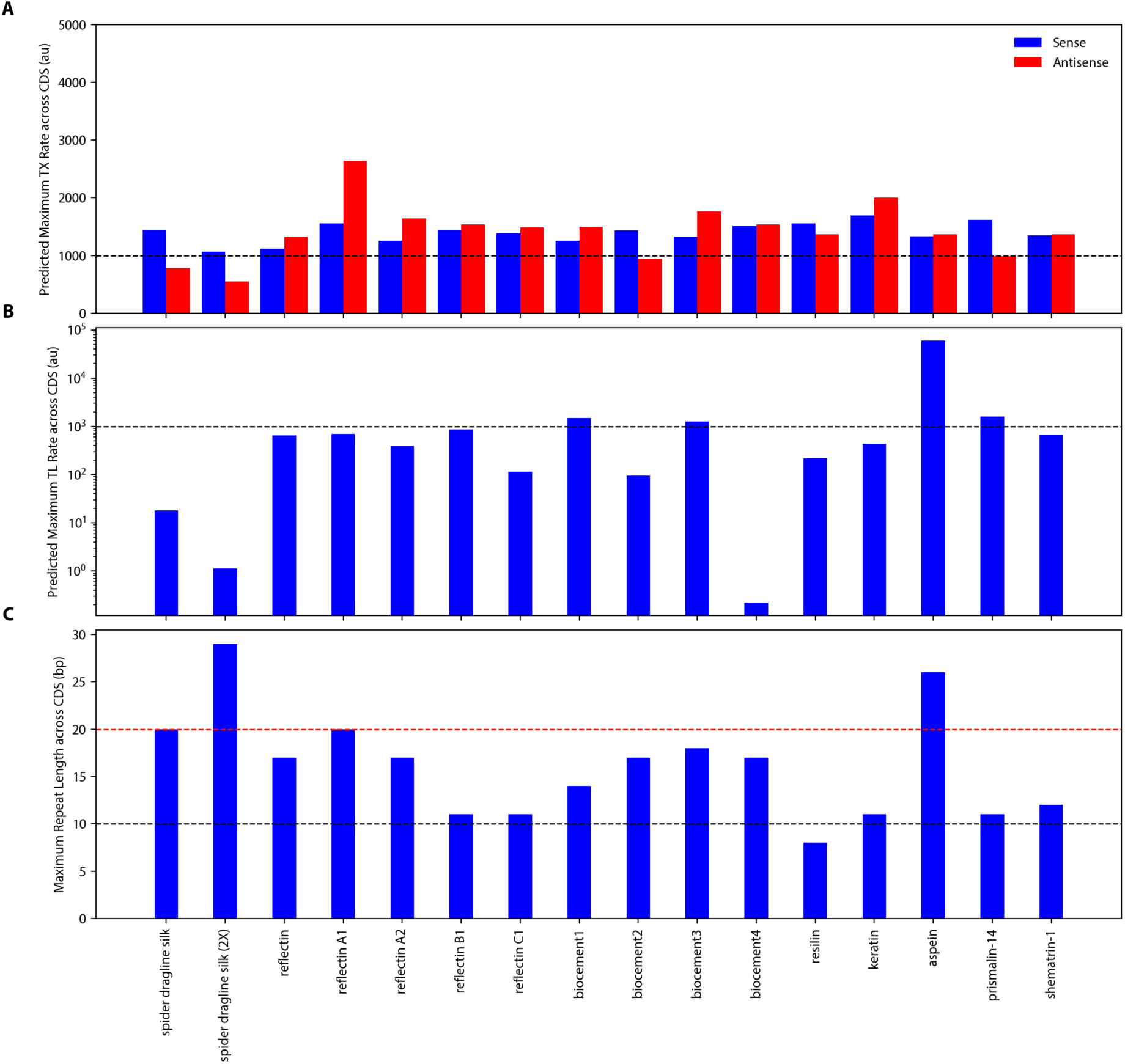
Structural protein CDS optimization. After applying the CDS Calculator to 16 structural protein CDSs, the transcription initiation rates, translation initiation rates, and maximum repeat lengths were calculated across each CDS sequence. **A.** The maximum values of the predicted transcription initiation rates in the (blue) forward/sense and (red) reverse/antisense directions are shown for each CDS as compared to the (black, dashed line) transcription rate of a weak promoter (1000 au). **B.** The maximum values of the predicted translation initiation rates across the optimized CDS sequences are shown as compared to the (black/dashed line) translation rate of a weakly translated internal ORF (1000 au). **C.** The maximum length of any repetitive DNA sequence is shown for each structural protein CDS as compared to a (black, dashed line) threshold repeat length of 10 bp and a (red, dashed line) repeat length of 20 bp, which increases the homologous recombination rate.

We then applied the RBS Calculator to design three ribosome binding site sequences for each CDS sequence, varying their translation initiation rates from moderate to very high levels (5000 to 300000 au on the RBS Calculator v2.1 scale). Transcription was driven by an engineered T7 promoter with a downstream lacO site for IPTG inducible expression. See **Supplementary Data 3** for the part compositions of all plasmids.

The extreme repetitiveness of many of these structural proteins makes them ideal examples to illustrate how applying multiple design rules can generate trade-offs. Trade-offs will occur when overlapping design rules (at their baseline thresholds) eliminate all possible sequence solutions. During sequence optimization, as design complexity increases, the adaptive changes in the design rules’ thresholds will favor some design criteria over others. For example, when all design rules are active, the CDS Calculator designed spider dragline silk CDSs that had no cryptic internal promoters or translated open reading frames (favoring these design rules) with maximum repeat lengths of 20 and 29 for the baseline and 2X versions, respectively. These trade-offs can be mitigated by adjusting the favored design rules’ baseline thresholds or deactivating some design rules altogether. For example, when we keep the “Minimize Repetitive DNA” rule activated and deactivate all other rules, we eliminate any possible trade-offs between design rules, resulting in maximum repeat lengths of only 15 for both versions, but at the cost of other design metrics; the resulting CDSs contained internal transcriptional start sites with maximum rates of 7000 to 9000 au. Overall, we can control the function of the designed CDSs by adjusting the design rules according to our desired application. In this scenario, the spider dragline silk CDSs did not have any appreciable translation of internal ORFs and any aberrantly produced mRNA isoforms would not be highly translated. However, in other examples, the transcription and translation of undesired mRNA isoforms would increase metabolic burden and provide a greater evolutionary push towards mutating the genetic system.

### Genetic Systems Builder: Design of Oligopools for Massively Parallel, Low-cost DNA Assembly

Massively parallel, low-cost DNA assembly is now possible by combining the development of many-fragment Golden Gate assembly with the low-cost economy-of-scale of oligopool synthesis. The DNA assembly workflow consists of carrying out PCR on a designed oligopool template using designed PCR primers that selectively amplifies a mixture of DNA fragments. After DNA purification, the DNA fragment mixture is then assembled into circular plasmid using Golden Gate assembly (**Figure 1B**). Importantly, there is no need for a researcher to purposefully introduce or pre-arrange overhangs within their assembled constructs. Instead, overhang selection is carried out at the algorithmic design stage, utilizing the overhangs already present inside each construct. Accordingly, to enable high-throughput DNA assembly using this approach, we developed and validated a novel algorithm (the Genetic Systems Builder) that designs the oligonucleotide sequences in the oligopool and the PCR primer pairs to maximize the efficiency of the many DNA assembly reactions. The algorithm also enables multi-level, one-pot library, and combinatorial assemblies using the same workflow.

The inputs into the Genetic Systems Builder algorithm include (**Figure 4A**): (i) the target melting temperature of all the PCR primers; (ii) the maximum length of the oligonucleotides in the oligopool; (iii) the Type IIS enzymes used for level 1, level 2, or combinatorial assembly, either BsaI, BbsI, Esp3I, BsmBI, or SapI; and (iv) either linear or circular assembly for each construct, where if linear assembly is selected, the terminal overhangs must be specified. The key algorithm outputs are: (i) the designed PCR primer sequences in plate format; (ii) the PCR primers’ melting temperatures; (iii) the designed oligonucleotide sequences; and (iv) the predicted efficiencies of each DNA assembly.

**Figure 4.**
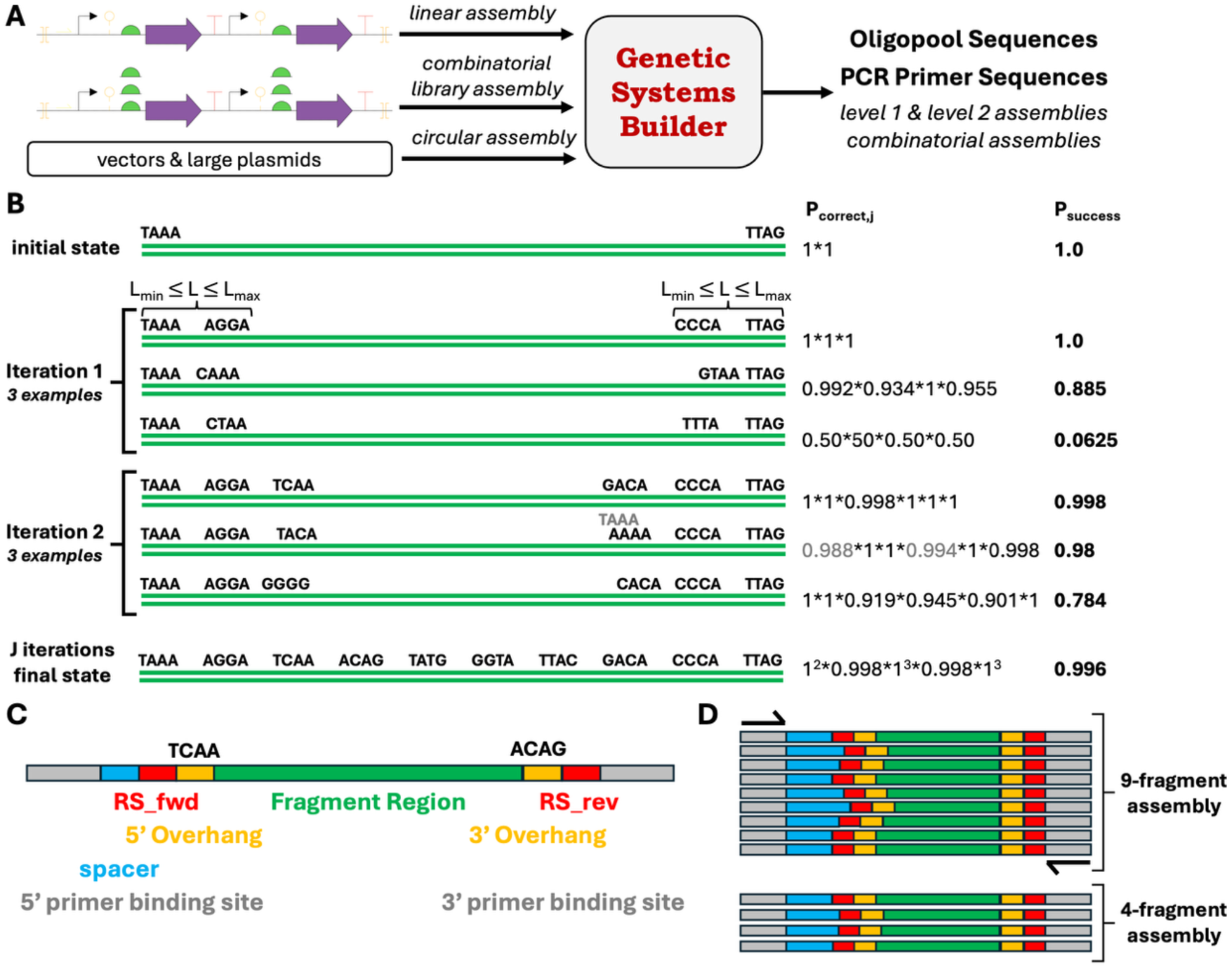
The Genetic Systems Builder. A dynamic programming algorithm designs oligonucleotide and PCR primer sequences to build many plasmids from a single oligopool. **A**. The algorithm supports linear assemblies into a vector, circular assemblies, and one-pot combinatorial assemblies from oligonucleotide variant libraries. **B.** The algorithmic procedure is shown for a linear assembly, using a beam width of 3, alongside the calculated probability of per-overhang and overall assembly. The algorithm also takes into account the presence of shifted overhangs that alters assembly probabilities (gray labels). **C.** The oligonucleotide sequences are designed after the optimal overhang sets are identified for each assembly, using the desired Type IIS restriction sites (RS). PCR primer binding sites are designed for highly specific amplification of oligonucleotide sub-pools. **D.** All oligonucleotides are synthesized in a single oligopool to maximize the molecular economy of scale.

The Genetic Systems Builder algorithm carries out several actions: (i) enumeration of the allowable overhang positions within the construct; (ii) identification of the overhangs and overhang positions that maximize overall ligation fidelity; (iii) design of PCR primer sequences that have optimal selectivity & amplification criteria; and (iv) the final design of the oligonucleotide sequences, incorporating the construct region, the flanking sequences (overhangs, assembly enzyme cut sites, primer binding sites), and designed padding sequences for uniform oligo lengths. The first two steps are carried out using dynamic programming optimization with beam search, while PCR primer and padding design are carried out using a sieving approach with several constraints (similar to our Oligopool Calculator algorithm^25^).

Dynamic programming with beam search is an iterative algorithm that begins with a decision tree, enumerates the next allowable decisions, ranks them using a scoring function, and retains a limited number of the top-scoring decision trees. The highest-scored solution is returned after completing the tree. The number of decision trees being evaluated in parallel is called the beam width ***W***. Here, the allowable decisions are the selected overhangs at specific positions in the construct. The decision tree is the set of overhangs selected to assemble the construct using Golden Gate assembly. The scoring function predicts overall assembly efficiency, taking into account the entire decision tree (all of the selected overhangs). During the overhang enumeration step, we exclude overhangs that were previously selected in the decision tree, overhang positions that would make the oligonucleotide lengths fall outside our allowable length range, and overhang positions located inside regulatory elements (for example, promoters and RBSs).

To predict assembly efficiency, we utilize the recently measured ligation frequencies of all pairs of overhang junctions for the most commonly used Type IIS assembly enzymes^34^. For all overhang pairs in each decision tree, we tally: (i) the frequency of correct assembly between the desired overhangs; (ii) the frequency of incorrect assembly between mis-paired overhangs, including ligations between two identical fragments and the reverse-complements of these overhangs; (iii) the frequency of the Type IIS enzyme mis-cutting the DNA (called slippage) and shifting the overhang and location by +1 or -1 bp; and (iv) the frequency of an incorrect assembly between the shifted overhang and other overhangs. The probability of a successful assembly ***P_i_*** at junction ***i*** is the ratio of correct assembly frequencies divided by the sum of correct and incorrect assembly frequencies. The overall assembly efficiency is the product of ***P_i_*** across all junctions. To estimate the frequency of Type IIS enzyme mis-cutting, we use the previously measured frequencies of enzyme mis-cutting for 14 restriction endonucleases^35^, which varied from 1.1% to 53.6%, as upper-bound limits as these characterized enzymes are not commonly used for DNA assembly. For BsaI, BbsI, Esp3I, BsmBI, and SapI, we use a probability of enzyme mis-cutting of 0.1% (an order-of-magnitude estimate), which will sufficiently penalize the use of shifted overhangs that produce undesired ligation reactions.

For long constructs with many overhang choices (more than what is needed for perfect assembly efficiency), we introduce ways to reduce overall computational runtime, while not sacrificing optimality. We eliminate the use of overhangs that have average ligation fidelities below a threshold, ***L_min_***. We also deterministically step over every ***N*** allowed overhang positions, using an adjustable step parameter ***N***. By adjusting the beam width ***W***, the minimum ligation fidelity ***L_min_***, and the step parameter ***N***, we can scale the computational intensity of the dynamic programming algorithm. The parameters for exhaustive enumeration of all possible choices are (***W*** = infinite; ***L_min_*** = 0; ***N*** = 1). Our default parameters suitable for most constructs are (***W*** = 2; ***L_min_*** = 0.60 to 0.85; ***N*** = 1 to 6). If no solution is found using the default parameters, our fall-back parameters are (***W*** = 3; ***L_min_*** = 0.60; ***N*** = 1), which was found to always return a solution on all tested sequences at the cost of greater computational runtime.

In **Supplementary Data 1**, we report the model-predicted ligation fidelities and computational runtimes of the Genetic Systems Builder algorithm across example sequences with varied lengths and complexities, including 206 plasmids expressing structural proteins, 44 vectors (up to 5600 bp), 3 phage genomes (up to 11200 bp), and 738 random sequences (with varying lengths from 3000 to 13000 bp). The Genetic Systems Builder required about 0.70 seconds (on average) per assembly with a maximum of 56 seconds.

After the algorithm is run and the designed DNA materials are synthesized, the experimental workflow is carried out using the protocol found in **Supplementary Data 2**. The protocol is fairly routine for PCRs and Golden Gate assemblies, though it is important to reduce the number of PCR cycles to maintain uniform amplification. We also carry out a heat soak on the PCR product to unfold intramolecular DNA structures prior to Golden Gate assembly.

### Assembly of plasmids using the Genetic Systems Builder workflow

We evaluated the performance of the Genetic Systems Builder algorithm by building 240 plasmids: 39 that express structural proteins with varied ribosome binding sites (up to 2000 bp inserts); 1 plasmid expressing mRFP1 as a positive control; 44 Golden Gate compatible vectors with varied inducible expression systems and selection markers (up to 5600 bp circular assemblies); and 156 plasmids expressing talin structural protein variants (up to 1700 bp inserts) (**Figure 5A**). Plasmids were constructed across 3 rounds, where the results from each round informed algorithmic improvements, culminating in a “v1.0” of the Genetic Systems Builder. Rounds 1 and 2 used routine transformation, colony-picking, and nanopore whole-plasmid sequencing to identify mutations and validate assembly success. Round 3 used pooled nanopore sequencing on mixed assembly product to directly measure assembly efficiency prior to any transformation or isogenic isolation. Both approaches leveraged our mapping algorithm (Auto Align) to automatically map nanopore reads to all reference sequences. We measured key metrics across the workflow, including PCR product concentrations, assembly efficiencies, and observed mutations.

**Figure. 5.**
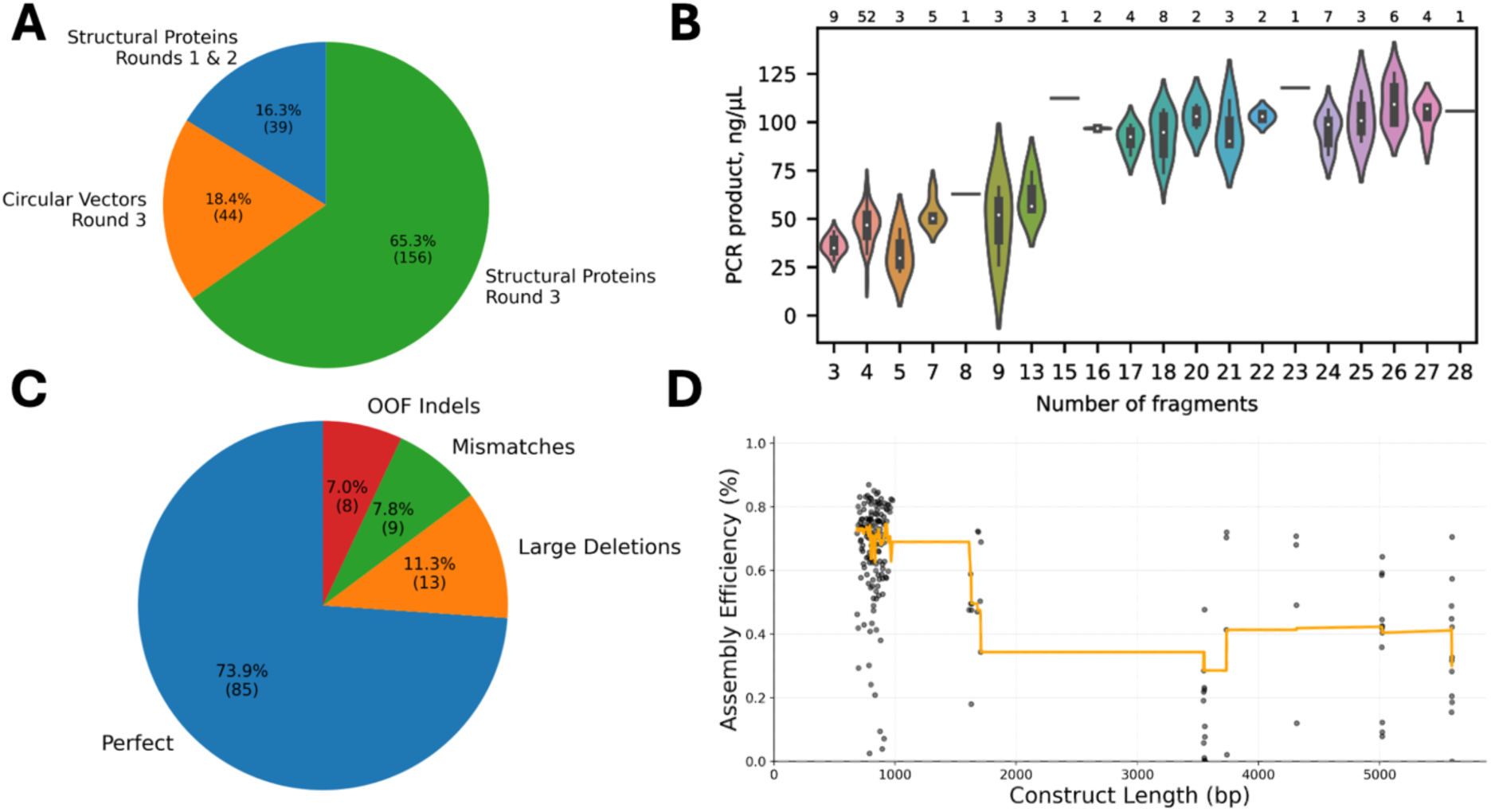
Stress testing DNA assembly by building structural protein expression plasmids. **A.** An overview of the numbers of plasmids produced across all assembly reactions. **B.** PCR product concentrations for 67 assembly reactions compared to the number of DNA fragments amplified in each PCR. The number of assembly reactions with the same number of DNA fragments is shown at the top. **C.** The outcomes of assemblies in rounds 1 and 2 (excluding aspein and dragline spider silk 2x), showing the number of sequenced, isoclonal plasmids that contained either perfect sequences, out-of-frame (OOF) insertion-deletions (indel) mutations, mismatch mutations, or large deletions. **D.** Measured efficiencies from pooled nanopore sequencing of Round 3 assemblies, compared to their construct lengths (black dots). The median assembly efficiency across each construct length is shown (orange line).

We first found that PCR product concentrations were approximately proportional to the number of fragments produced in the first step of the workflow (Pearson R = 0.644, p = 0.0175, N = 67) (**Figure 5B**). A proportional relationship is expected when each primer pair is capable of selectively amplifying the target set of DNA fragments needed for each downstream many-fragment Golden Gate assembly. However, we do observe a concentration plateau after about 20 fragments, suggesting that these lower-cycle PCR reactions or the downstream DNA purification can become yield-limiting.

In rounds 1 and 2, we utilized the Genetic Systems Builder’s library assembly mode to build 3 versions of the structural protein expression plasmids in each well (containing the ribosome binding site variants) along with the mRFP1 positive control in a separate well. We picked & sequenced 129 colonies and identified any mutations in their plasmids, categorizing them as either single mismatches, insertion-deletions that cause out-of-frame shifts, or large deletions (**Figure 5C**). Large deletions are caused by incorrect ligations during DNA assembly, which is important to assessing the Genetic Systems Builder’s performance. Whereas, single mismatch mutations are predominantly caused by errors during oligonucleotide synthesis, while insertion-deletion mutations can be generated during DNA repair after transformation.

Overall, we built fully correct plasmids for 14/16 of the structural proteins (88% success rate), each having one or more ribosome binding variants that vary their expression levels (36 plasmids total). The two failures were aspein (N = 9) and the longer version of spider dragline silk (N = 5). We hypothesize that leaky expression of these proteins (even without induction) created significant toxicity that precluded the plasmids’ replication inside *E. coli* cells. Across the remaining 115 plasmids, 74% were fully correct, 11% had large deletions, 8% contained mismatches, and 7% had out-of-frame insertion-deletions (**Supplementary Data 3**).

In round 3, we used the Genetic Systems Builder to assemble 200 plasmids, utilizing the circular assembly mode to build 44 vectors and the linear-into-vector assembly mode to build 156 Talin expression plasmids. Vectors varied in length from 3500 to 5600 bp (15 to 24 assembled fragments) and contained over 20 genetic parts, including a ColE1 origin, expression of LacI for inducible transcriptional control, expression of chloramphenicol acetyltransferase for CmR selection, homology arms for genome integration at desired sites, high-fidelity junctions for Golden Gate assemblies, and several terminators to insulate expression; multiple versions were constructed, for example, varying the translation rates of LacI and CmR, for different applications. The Talin expression cassettes varied in length from 700 to 1700 bp (3 to 8 assembled fragments) and contained over 10 genetic parts to inducibly express variants of the Talin structural protein, utilizing different inducible promoters, purification and protease tags, and ribosome binding sites to vary Talin expression levels. PCRs, DNA purification, and Golden Gate assemblies were carried out in 96-well plate format for streamlined operation.

Across the 200 DNA assemblies, we combined together assembly product prior to any DNA purification, carried out pooled nanopore sequencing, and mapped long reads to reference sequences using the Auto Align mapper, which assigns each read to the closest matching reference sequence. There were 994500 mapped reads (median N50 of 4156 bp after filtering) with a median of 5542 reads per reference sequence. We then calculated the number of mapped reads ***NR_i_*** that have matching sequence at each position ***i*** across the reference sequence. The assembly efficiency is the ratio of the minimum number of matched positions over the maximum number of matched positions, while correcting for a nanopore base calling error ***Eps*** of about 0.06^36^, or ***E/*** = min(***NR_i_***) / max(***NR_i_***) / (1 – ***Eps***). The assembly efficiency for the 44 large vectors was 33%, while the assembly efficiency for the Talin expression cassettes was 65% (**Supplementary Data 4**). Increasing the number of DNA fragments in the assembly lowered its efficiency; a simple linear fit (R^2^ = 0.31, p = 1.5x10^-17^) shows that each additional DNA fragment decreased efficiency by about 1.85%, starting from a baseline of 72% (**Figure 5D**).

### Low Material Costs for High-throughput DNA Assembly using the Genetic Systems Builder

The costs of DNA synthesis, assembly, and sequencing are a critical factor to implementing a high-throughput genetic systems engineering pipeline. Here, we tally the costs of the consumable reagents needed to build many plasmids from a single oligopool and verify their sequences using pooled nanopore sequencing, while using the Genetic Systems Builder algorithm for oligopool and PCR design. These costs include the thermostable DNA polymerase, PCR primers, oligopool, T4 DNA ligase, Type IIS restriction enzymes, DNA purification kits, microtiter plates, and pooled nanopore sequencing, purchased from New England Biolabs, Integrated DNA Technologies, Twist Biosciences, Zymo Research, and Plasmidsaurus. While material costs do vary across manufacturers and grow with inflation, it is important to carry out such analyses to provide a checkpoint in time and to enable systematic comparisons across techniques.

We find that the material costs of the Genetic Systems Builder approach is about $0.0276 to $0.005/ bp, which is about 2.5 to 24-fold lower than current commercial service providers ($0.07 to $0.12/bp), depending on the sequence complexity, scale, and insert size (see **Supplementary Data 5** for a cost calculator). Specifically, for 1200 bp inserts, building 100 plasmids costs about $33/plasmid ($0.0276/bp), while building 1000 plasmids costs about $19/plasmid ($0.0156/bp). For 4800 bp inserts, building 100 plasmids costs about $54/plasmid ($0.0112/bp), while building 1000 plasmids costs about $26/plasmid ($0.0053/bp). Overall, there is a log-linear relationship between the number of assemblies per oligopool and the per-unit assembly costs. We call this relationship a *molecular economy of scale*, which incentivizes the construction of many genetic systems at the same time to drive down costs. In practice, this often means that multiple projects worth of oligopool DNA should be ordered at the same time. The material cost calculation excludes labor and other operating costs; a more direct cost comparison would depend on many external factors.

Based on the cost itemization, the predominant material cost is the oligopool itself (50-76% of each assembly) with the PCR primers and T4 DNA ligase as the two main secondary costs (5-15% of each assembly). If additional cost savings are needed, it is possible to design a set of universal PCR primers that can be re-used across workflows using different oligopools. Re-using PCR primers will save up to $4 per assembly, but will add additional storage and logistical requirements.

### Quantifying Single-cell Protein Expression Using a Translationally Coupled Biosensor

Next, we developed an intracellular biosensor that measures the expression level of a protein-of-interest, without the use of fusion tags, in order to overcome the testing bottleneck. To do this, we leveraged the mechanism of translational coupling and ribosome re-initiation to generate a proxy fluorescence signal that is predicted to be perfectly proportional to the protein-of-interest’s expression level. Specifically, when introducing a protein-of-interest CDS into an IPTG-inducible T7 expression system, we utilize an overlapping ATGA stop-start junction with a downstream mCherry fluorescent protein reporter (**Figure 6A**). When ribosomes translate the upstream coding sequence and produce the protein-of-interest, there is a small probability that the ribosome dissociates and immediately re-initiates translation of the downstream mCherry fluorescent reporter (**Figure 6B**). Based on a biophysical model of translational coupling^13^, the mCherry fluorescence level is predicted to be proportional to the expression of the upstream protein-of-interest.

**Figure. 6.**
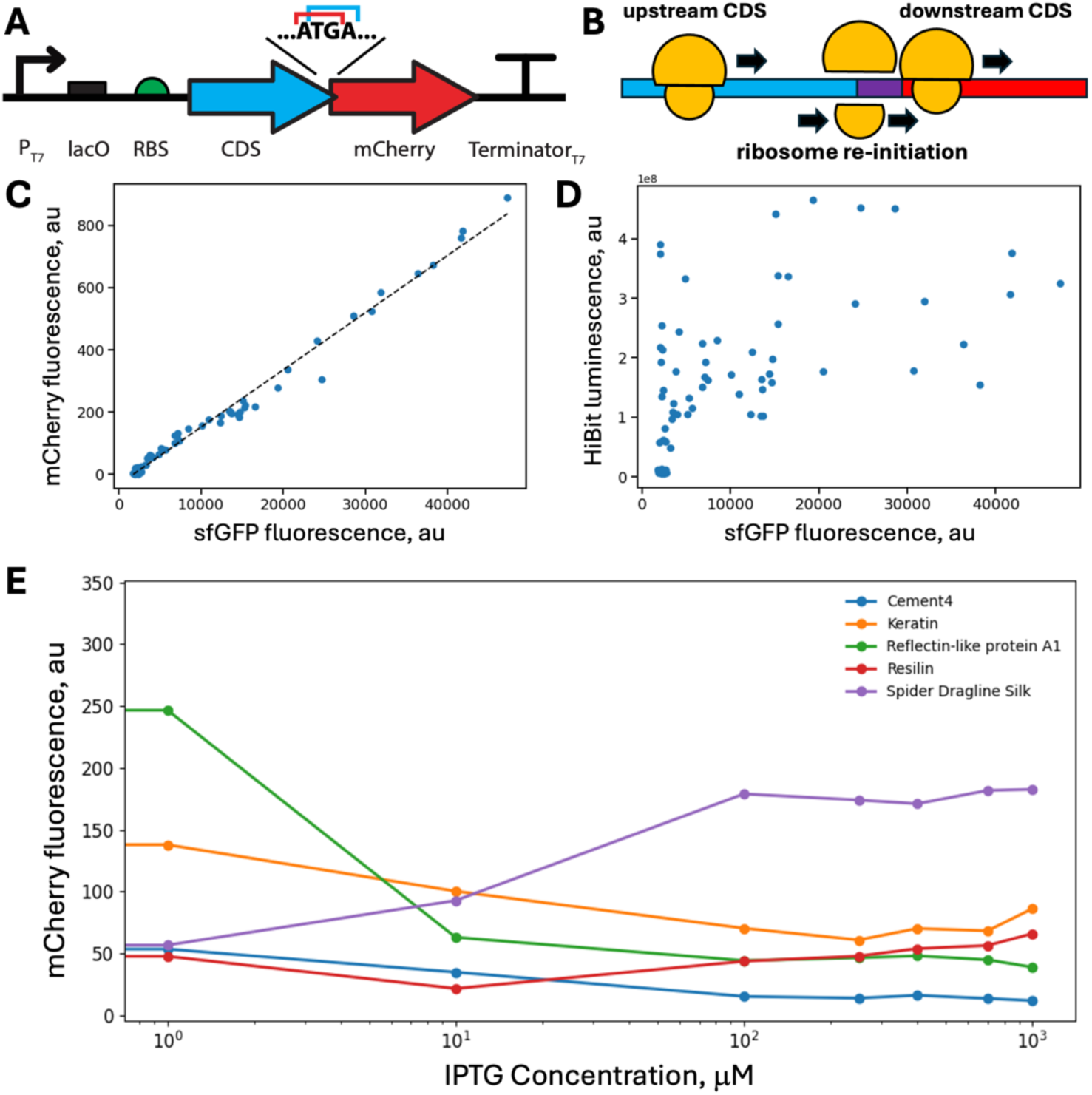
Translationally coupled biosensor for single-cell protein expression characterization. **A.** Schematic of the IPTG-inducible T7 expression vector, showing overlapping start and stop codons (ATGA) between the protein-of-interest CDS and the translationally coupled mCherry biosensor. **B.** Ribosome re-initiation occurs when the 70S ribosome dissociates at a stop codon and the 30S ribosomal subunit rebinds at the nearby start codon, translating the downstream CDS in proportion to the upstream CDS. **C.** Measured fluorescence levels from a sfGFP protein-of-interest in comparison to the translationally coupled mCherry biosensor fluorescence levels. (Pearson r = 0.993, p = 1.5x10^-74^, points are the average of 2 biological replicates) **D.** Measured fluorescence levels from a sfGFP protein-of-interest in comparison to luminescence levels from a HiBit tag. (Pearson r = 0.592, p = 7.4x10^-9^, points are the average of 2 biological replicates) **E.** Expression level measurements for several structural proteins, showing how biosensor fluorescence levels change in response to increasing IPTG inducer concentrations (points are the average of N = 2 biological replicates).

We validated the performance of this tag-free biosensor by inserting sfGFP as the protein-of-interest and systematically varying its expression level using a combination of IPTG induction and different RBS sequence variants. The RBS variants were designed to vary translation initiation rates from 17000 to 180000 on the RBS Calculator proportional scale. sfGFP fluorescence emission is spectrally orthogonal to mCherry excitation. We also tagged the sfGFP with a C-terminal HiBit tag to carry out a direct comparison with the conventional HiBit luminescence assay. From identical split cultures, sfGFP and mCherry fluorescence were measured via flow cytometry, while HiBit luminescence levels were quantified in parallel inside microtiter plates.

Flow cytometry revealed an extremely high correlation between the biosensor mCherry fluorescence level and the protein-of-interest sfGFP fluorescence level (Pearson r = 0.993; two-tailed T-test p = 1.5x10^-74^; **Figure 6C**). HiBit luminescence measurements from the same, split cultures were also correlated with sfGFP fluorescence levels, but at a much lower level (Pearson r = 0.59; two-tailed T-test p = 7.4x10^-9^ **Figure 6D**). Unlike the intracellular biosensor measurements, small differences in volumes and timing can result in significant differences in HiBit luminescence levels. Raw fluorescence and luminescence levels are provided in **Supplementary Data 6**.

After validation and calibration, we applied the translational coupling biosensor to measure the expression levels of several structural proteins, including spider dragline silk, keratin, reflectin-like protein A1, resilin, and three bio-cements (Cement1, Cement3, Cement4). As we varied transcription rates by increasing IPTG induction, we recorded distinct CDS-specific responses that suggest protein-dependent global effects (**Figure 6E**). For spider dragline silk, an increase in transcription rate led to an overall increase in protein expression (as expected) up until a plateau region is reached. However, for the proteins reflectin, cement4, and keratin, we found that increasing transcription rates led to a reduction in protein expression, likely due to toxicity effects. Resilin expression remained relatively flat across these conditions. These high-throughput measurements quantify what is commonly observed as a “less is more” behavior for structural proteins, where slowing expression down can actually lead to higher protein levels.

We then investigated how changing both transcription and translation rates could alter protein expression levels across a cell population. We measured the single-cell biosensor fluorescence levels when expressing Cement1, using RBS variants with a low, medium, or high translation initiation rate. With a low or medium translation initiation rate, the Cement1 protein followed a “more is more” pattern where higher transcription rates (more IPTG induction) resulted in higher average expression levels with a single peaked, Gaussian-like distribution of single-cell expression levels (**Figure 7AB**). The “medium” RBS translation rate yielded the highest amounts of Cement1 protein. However, once the translation initiation rate reached a critically high point, the Cement1 expression levels remained low even as transcription rates were increased; the single-cell expression levels showed a dispersed and bimodal distribution with a large number of cells that do not express Cement1 protein. This behavior suggests that the high translation initiation rate may have exceeded the ribosome’s translation elongation rate, leading to ribosome bottlenecking and sequestration.

**Figure. 7.**
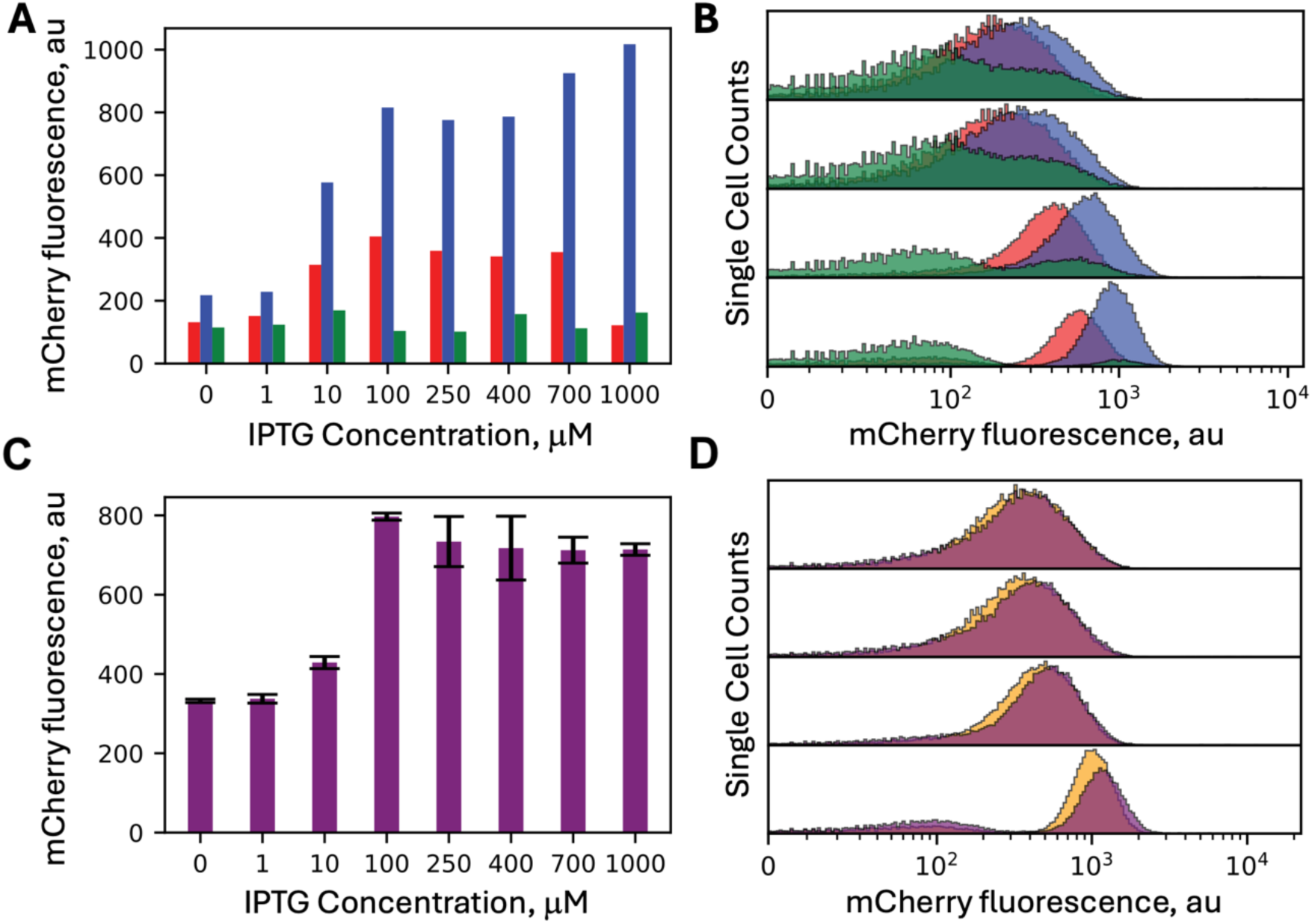
Multimodal protein expression levels in heterogeneous cell populations. **A.** Expression level measurements for the Cement1 structural protein, showing how average biosensor fluorescence levels change as IPTG inducer concentration is increased across expression constructs utilizing different ribosome binding site variants. Bars show RBS variants with either (red) low translation rates, (blue) medium translation rates, or (green) high translation rates. **B.** Overlaid single-cell fluorescence distributions showing changes in Cement1 expression level bimodality when induced with (top to bottom) 0 µM, 1 µM, 10 µM, or 100 µM IPTG. Colors indicate the RBS translation rates. **C.** Expression level measurements for the Cement3 structural protein, showing how average biosensor fluorescence levels change as IPTG inducer concentration is increased, using a RBS variant with a high translation rate. **D.** Overlaid single-cell fluorescence distributions showing changes in Cement3 expression when induced with (top to bottom) 0 µM, 1 µM, 10 µM, or 100 µM IPTG. Colors indicate expression differences between two isogenic cultures.

We then applied the translationally coupled biosensor to investigate the differences in protein expression level across isogenic colonies, using the Cement3 as a representative example. We selected isogenic strains, cultured them identically, and measured their single-cell Cement3 expression levels across a range of transcription rates (**Figure 7CD**). We found only small differences in Cement3 expression (average CV less than 10%) across biological replicates, providing a valuable way to quantify non-genetic differences in expression across cell populations. Overall, the translationally coupled biosensor provided a quantitative assessment of single-cell expression using a high-throughput and tag-free assay, enabling systematic comparisons of system parameters that control protein expression levels.

## DISCUSSION

We developed an integrated, high-throughput Design-Build-Test pipeline to overcome key bottlenecks in synthetic biology and accelerate genetic systems engineering. We combined multi-objective & model-based sequence optimization, a scalable & low-cost DNA assembly technique, pooled DNA sequencing, and high-throughput quantitative measurements of single-cell protein expression to enable efficient construction and characterization of complex genetic systems and libraries. We stress-tested this pipeline on a collection of structural proteins that are particularly challenging to express, due to their highly repetitive amino acid sequences and biased amino acid frequencies. Our pipeline designed, built, and tested genetic systems that express many structural proteins, while reducing the DNA assembly costs by up to 24-fold and quantifying the heterogeneity in single-cell protein expression levels. It is important to stress-test these pipelines on challenging systems to determine their effectiveness and robustness across real-world applications.

A key theme of our integrated pipeline is that decisions made in the design phase will control the success rate during the build and test phases. In this work, we introduce sequence optimization rules that improve build success (“design for build”) and enable high-throughput measurements (“design for test”), while controlling transcription rates and translation rates (“design for function”). These design principles are widely used in other engineering disciplines, particularly in microprocessor engineering, but are not widely used in synthetic biology. Specifically, we introduce new optimization rules to improve genetic stability and evolutionary robustness, lower the production of undesired RNAs and proteins (lowering metabolic burden and toxicity), and directly couple the protein-of-interest’s expression level to an easily measured proxy that is quantified at the single-cell level.

The CDS Calculator algorithm applies a highly novel approach to the common “codon optimization” problem. Current approaches mainly focus on varying synonymous codons to increase translation elongation rates, while not accounting for other sequence motifs or kinetic rates that control genetic system stability and function. Instead, our pipeline applies 15 design rules to control transcription initiation & elongation rates, translation initiation & elongation rates, mRNA stability, and RNAP and ribosomal pause sequences. Our design rules also remove undesired sequences that cause failures at the build and test phases, including repetitive sequences, mutational hotspot sequences, internal promoters, highly translated internal ORFs, and internal terminators. However, as more design rules are added to the multi-objective sequence optimization algorithm, we do find that certain CDS sequences can not be designed to satisfy all rules simultaneously, creating trade-offs during optimization. In this scenario, we must choose which design rules have the most impact on genetic system function, for example, prioritizing the removal of highly translated ORFs over the removal of internal promoters. A key next step to genetic systems engineering is to develop system-wide models that predict how these design choices control higher-order behaviors (e.g. metabolic burden), enabling improved prioritization of the design rules.

The Genetic Systems Builder algorithm extends these design principles into the build step by utilizing dynamic programming to rapidly identify optimal overhang sets for many-fragment Golden Gate assemblies, while designing highly selective primers for multiplexed PCRs across many thousands of oligonucleotides. We show that the Genetic Systems Builder achieves very high first-pass assembly efficiencies (65-75%, up to 2000 bp inserts) with the ability to build very large plasmids (33% efficiency, up to 5600 bp inserts). Importantly, unlike other approaches, we do not use any low-throughput or hand-heavy steps, such as gel electrophoresis, that would ultimately constrict throughput. The Genetic Systems Builder also designs multi-level hierarchical and combinatorial Golden Gate assemblies, automatically identifying the level 2 overhangs and adding the secondary flanking Type IIS cut sites within the oligonucleotides. Beyond the plasmids constructed in this work, several new researchers have been trained to use this approach to build over 1000 plasmids (900 to 2700 bp inserts), indicating its robustness and reliability. Achieving high-throughput and robust DNA assembly is particularly important as laboratories utilize more automated liquid handling robots to streamline operations.

Oligopools have been increasingly utilized for library-based cloning, site-saturation mutagenesis, sgRNA guide libraries, and DNA fragment assembly^18, 26, 28^. Over the same period, the efficiency of the Golden Gate assembly method has been optimized to improve its many-fragment efficiency^24^. The Genetic Systems Builder further improves on these methods by identifying Type IIS cleavage slippage as a key reason why mis-assemblies can occur by introducing shifted overhang sequences that happen to ligate to another overhang with an appreciable probability. We introduce a novel use of dynamic programming to identify optimal overhang sets, which directly takes into account Type IIS cleavage slippage and prevents the selection of ligation-prone shifted overhangs. Importantly, prior approaches have commonly used stochastic optimization to identify overhang sets, which can return different overhang sets for the same input DNA sequence. Such stochastic algorithms are easy to implement, but generate irreproducible results. Instead, the Genetic Systems Builder uses a deterministic algorithm that always returns the same, optimal overhang set for an input DNA sequence.

Pooled long-read sequencing has become the preferred method of assessing the correctness of assembled plasmids and plasmid libraries, due to its fast turnaround and low-cost. Here, we show that directly pooling DNA assembly product, before transformation and colony-picking, can provide effective quality control at a very low cost. For example, 1000 plasmid assemblies (5000 bp long) combined in one pool can be sequenced with 200-fold coverage from 1 Gbp of nanopore sequencing (costing about $500), yielding a cost of about $0.50 / plasmid. The cost benefits are even greater when sequencing combinatorial plasmid libraries with many thousands of pooled sequences. However, the presence of so many sequences together in one pool places a greater burden on the computational read mapping process. Existing read mapping algorithms (e.g. minimap2^37^) were not developed to map read sequences to thousands of long reference sequences. The algorithmic challenge is that many of these reference sequences can be highly similar with overlapping kmer distributions, which leads to long runtimes.

To resolve this challenge, we apply a kmer compactification strategy, which removes over-represented kmers from the reference sequences, carries out read mapping across the compactified references, and then rapidly reconstructs alignments between the mapped reads and original reference sequences. These improvements were introduced into a modified minimap2 repository with options to activate kmer compactification as needed. The benefit of this strategy generally increases with the number of references. For example, 1.3 million long reads were mapped to 50710 reference sequences in 59 minutes (372 reads per second), originating from a combinatorial plasmid library. The same dataset would require 8.5 days of runtime using the original minimap2 algorithm with a very slow mapping rate of only 1.8 reads per second. With so many reference sequences, the kmer compatification strategy reduced the computational runtime by 206-fold, removing a significant bottleneck in the data analysis pipeline. In this work, 12.2 million unfiltered reads were mapped to 200 reference sequences in 34 minutes (6000 reads per second), yielding a 1.5-fold speed-up.

The “test” stage remains one of the least scalable aspects of current Design–Build–Test workflows. Protein expression measurements are often dependent on fusion tags, destructive lysis-based assays, or population-averaged measurements that obscure instability and phenotypic heterogeneity. Our translational coupling biosensor addresses this limitation by linking expression of a fluorescent reporter directly to translation of the upstream protein-of-interest without modifying the target protein itself. Translational coupling and ribosome reinitiation mechanisms have previously been characterized in bacterial and archaeal systems^13, 38^. Here, we leverage this mechanism as a scalable quantitative biosensor for high-throughput protein characterization. Compared to luminescence-based assays such as HiBit, the translational coupling biosensor substantially reduces sample preparation requirements and reagent costs while remaining compatible with flow cytometry and automated fluorescence measurements. The ability to quantify single-cell distributions rather than population averages proved particularly valuable for identifying heterogeneous expression states and bistable-like transitions that would otherwise remain hidden.

The observed multimodal expression distributions suggest that expression burden and translational bottlenecking generate heterogeneity at the single-cell level, particularly when over-expressing heterologous proteins. The appearance of bimodal subpopulations at high transcription or translation initiation rates suggests that translational capacity becomes limiting once ribosome loading exceeds elongation throughput. Similar resource-allocation and burden-driven transitions have been reported in other genetic systems, including T7 expression systems and synthetic burden circuits^39-41^, although they are often difficult to detect using bulk measurements alone. Our results highlight the importance of incorporating single-cell phenotyping into high-throughput genetic system optimization workflows.

The techno-economic implications of this workflow are also significant. Oligopool-enabled assembly introduces a molecular economy of scale in which per-construct costs decrease as library size increases. This contrasts sharply with conventional gene synthesis workflows, where costs scale approximately linearly with construct number and sequence length. Although additional sequencing and validation remain necessary, the reduction in material costs fundamentally changes the feasibility of large-scale experimental exploration. Importantly, this cost reduction is likely to become even more pronounced as oligopool synthesis lengths increase and synthesis fidelity improves.

While the presented pipeline has substantially expanded and accelerated synthetic biology workflows, there are still limitations. First, some structural proteins remain highly challenging to express, for example, due to non-specific interactions inside cells that interfere with essential processes. This pipeline makes it possible to screen large quantities of helper enzymes to facilitate protein folding, de-aggregation, and sequestration, which will increasingly be needed as larger protein complexes are introduced into heterologous hosts. Novel tag-free sensors for measuring *in vivo* protein folding, solubility, activity, and cell state can be incorporated into larger sensor arrays that provide a more holistic view of intracellular processes, expanding the “design for test” principle.

The pipeline software is available on GitHub at https://github.com/hsalis.

## METHODS

### Vector Design, DNA Assembly and Cloning

The pHMS2 vector (ColE1, CmR, LacI) was designed using the Promoter Calculator, RBS Calculator, and CDS Calculator to express chloramphenicol acetyltransferase and the LacI transcription factor at desired expression levels. The protein-of-interest expression cassette was designed with BsaI outie sites using 5’ TAAA and 3’ TTAG terminal overhangs, followed by an in-frame C-terminal HiBit tag, a stop-start translational coupling motif, and the mCherry CDS. Prior to the development of the Genetic Systems Builder, the vector was constructed via PCR amplification of four DNA fragments (gBlocks, Integrated DNA Technologies) and carrying out Gibson assembly (NEBuilder HiFi, New England Biolabs). Purified isoclonal plasmids were verified using nanopore sequencing (Plasmidsaurus). Vector sequences are available in **Supplementary Data 7**.

The Genetic Systems Builder workflow was applied to build the structural protein expression plasmids and the sfGFP expression plasmids for calibration of the translational coupling biosensor, using pHMS2 as the vector and the protocol in **Supplementary Data 2**. Plasmids were transformed into NEB Stable cells (E. coli MG1655 F’ *proA^+^B^+^ lacI^q^ Δ(lacZ)M15 zzf::Tn10 (*Tet^R^*)/Δ(ara-leu) 7697 araD139 fhuA ΔlacX74 galK16 galE15 e14- Φ80dlacZΔM15 recA1 relA1 endA1 nupG rpsL* (Str^R^) *rph spoT1 Δ(mrr-hsdRMS-mcrBC*), recovered using supplied outgrowth media (New England Biolabs), and plated on LB agar (20 μg/mL chloramphenicol) for isoclonal isolation. Cultures were carried out at either 37°C or 28°C, depending on the expression level and apparent toxicity of the protein-of-interest, using 300 RPM orbital shaking. Construction of circular vectors used the same procedure, except excluding the addition of the pHMS2 acceptor vector. Plasmid sequences were verified using nanopore sequencing (Plasmidsaurus) using either isoclonal or pooled sequencing. Consensus or raw read sequences were mapped to reference sequences using the Auto Align algorithm.

### Computational Design of Genetic Parts

The structural protein CDSs were designed using the CDS Calculator. By default, all design rules were activated, except where noted. BsaI and BbsI restriction sites were excluded. The targeted maximum repeat length was 10. The ribosome binding site sequences were designed using the RBS Calculator to systematically vary translation initiation rates across levels with targeted rates being either low (5000), medium (40000), or high (300000), including the structural protein CDS sequences as inputs. Actual predicted translation initiation rates varied across CDS sequences, depending on target rate feasibility.

### Protein Expression Measurements and Characterization

Structural protein and sfGFP expression plasmids were transformed into *E. coli* BL21(DE3) cells (New England Biolabs), plated, and cultured in LB media supplemented with 20 μg/mL chloramphenicol and varied IPTG concentrations (0 to 1000 μM) at 28°C with 300 RPM orbital shaking. Multiple successive cultures were carried out using 1:50 serial dilutions for a total culture period of 24 hours. Non-transformed *E. coli* BL21(DE3) cells were used as negative autofluorescence controls. OD_600_, sfGFP fluorescence, and mCherry fluorescence were recorded using a spectrophotometer (Tecan Spark) every 5 minutes. Flow cytometry (BD LSR Fortessa) was used to record single-cell sfGFP and mCherry fluorescence distributions of cell samples, using an ellipsoidal FSC and SSC gate that captured bacterial cells. Reported sfGFP and mCherry fluorescence levels were calculated by first subtracting the autofluorescence and then taking the arithmetic mean over replicates.

HiBit-based expression measurements were carried out using the Lytic HiBit assay (Promega). Cell samples were fixed by transferring to a PBS solution with 2 mg/mL kanamycin, which halts protein synthesis. The HiBit solution was prewarmed and combined with fixed cells using the manufacturer’s recommended dilution factor, followed by incubation at 24°C. Luminescence levels were recorded every 5 minutes (Tecan Spark). BL21(DE3) cells were used as the negative control to measure any autoluminescence. Reported luminescence levels are the endpoint luminescence levels with autoluminescence subtracted, then divided by cell biomass (corrected OD_600_), and taking the arithmetic mean over all replicates.

## ACKNOWLEDGEMENTS

This project was supported by funds from DefenseWerx and the National Science Foundation (DBI-2400302). The views and conclusions contained herein are those of the authors and should not be interpreted as necessarily representing the official policies or endorsements, either expressed or implied, of the U.S. Government.

